# An improved ATP FRET sensor for yeast shows heterogeneity during nutrient transitions

**DOI:** 10.1101/2019.12.12.874115

**Authors:** Dennis Botman, Johan H. van Heerden, Bas Teusink

**Author notes:** Correspondence should be addressed to Bas Teusink.

## Abstract

Adenosine 5-triphosphate (ATP) is the main free energy carrier in metabolism. In budding yeast, shifts to glucose-rich conditions cause dynamic changes in ATP levels, but it is unclear how heterogeneous these dynamics are at the single-cell level. Furthermore, pH also changes and affects readout of fluorescence-based biosensors for single-cell measurements. To measure ATP changes reliably in single yeast cells, we developed yAT1.03, an adapted version of the AT1.03 ATP biosensor, that is pH-insensitive. We show that pregrowth conditions largely affect ATP dynamics during transitions. Moreover, single-cell analyses showed a large variety in ATP responses, which implies large differences of glycolytic startup between individual cells. We found three clusters of dynamic responses, and show that a small subpopulation of wild type cells reached an imbalanced state during glycolytic startup, characterised by low ATP levels. These results confirm the need for new tools to study dynamic responses of individual cells in dynamic environments.

## Introduction

Adenosine 5-triphosphate (ATP) is one of the key players in cellular metabolism as it is the major Gibbs energy-carrier for most if not all species^1,2^. ATP is produced either by proton-gradient driven ATPases, or by substrate-level phosphorylation. The latter occurs in glycolysis, the central metabolic pathway in many organisms, including human. A key question is how pathway flux is regulated under dynamic conditions, which happens in nature, biotechnical processes^3–6^, but also in humans^7–9^. Given the task of glycolysis to produce ATP, it is not a surprise that the glycolytic flux is (also) regulated by ATP itself ^10,11^. *Saccharomyces cerevisiae* (or budding yeast) has been the focus of many studies of glycolysis; when it encounters an environmental change to glucose-rich conditions, glycolysis rapidly becomes active ^12–14^. For glycolysis to run, an initial investment of 2 ATP in the upper part (the conversion from glucose to fructose-1,6-bisphosphate) is needed, which is succeeded by a return of 4 ATP in the lower part (the conversion from glyceraldehyde-3-phosphate to pyruvate, in twofold, Fig. 1). When glucose is added, the initial and rapid use of ATP thus creates a (temporarily) imbalance between the two parts of glycolysis, resulting in a transient decrease of ATP levels^14–16^. The initial difference in flux between the two parts of glycolysis can recover, resulting in a balanced state. However, the upper glycolysis flux can also continue to exceed the lower glycolysis flux, resulting in an imbalanced state^16–18^.

**Figure 1.**
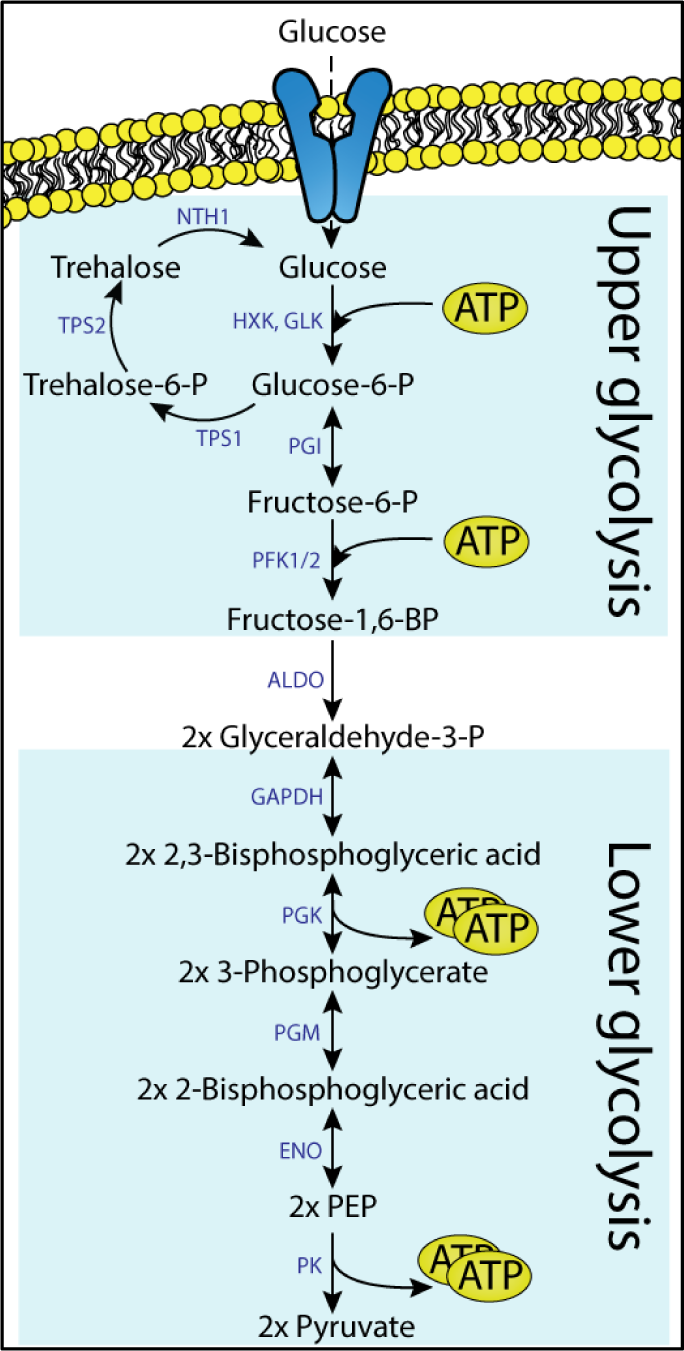
Schematic overview of glycolysis. Startup of glycolysis requires an investment of 2 ATP molecules in the upper part of glycolysis. The lower part of glycolysis yields 4 ATP afterwards.

These studies therefore suggested that startup of glycolysis can be highly variable, with a small fraction of cells dynamically ending up an imbalanced metabolic state^17^. This can be important for industrial processes in which metabolic subpopulations can affect industrial efficiency^3–6^. Such variability could also have implications for therapeutic efficiencies in human diseases^19–22^. These conclusions, however, were based on computational modelling and indirect evidence, in particular pH was used as an indirect readout of metabolism, not ATP itself. There is a need, therefore, to directly monitor ATP dynamics at the single cell level and establish a low ATP, imbalanced state. However, continuous measurements of ATP, and hence glycolytic startup dynamics, have not been studied at a single-cell level.

In recent years several fluorescence-based biosensors for *in vivo* monitoring of ATP in single cells have been developed, including the AT1.03 and the QUEEN sensor^23,24^. However, these sensors use fluorescent proteins (FPs) that are pH-sensitive in the physiological range where the intracellular pH of budding yeast operates^17,25–29^. In fact, the pH transiently drops exactly during glycolytic startup^17,30^. Moreover, the metabolic imbalanced state leads to an inability to maintain pH homeostasis, resulting in a significantly drop in intracellular pH, making the sensors unsuitable to study ATP dynamics during glycolytic startup. Therefore, we adapted the AT1.03 sensor to become less pH sensitive. This modified sensor, denoted yAT1.03, is pH-robust, and can be used to reliably detect single-cell ATP dynamics.

## Material and methods

### yAT1.03 construction

The ATP sensor AT1.03 and its inactive variant AT1.03^R122KR126K^ in the yeast expression vector pDRF1-GW were a gift from Wolf Frommer (Addgene plasmids #28003 and #28005, respectively). First, an EcoRI restriction site was removed by performing a PCR on AT1.03 and AT1.03^R122KR126K^ pDRF-GW using KOD polymerase (Merck-Millipore, Burlington, Massachusetts, USA) with the forward primer 5’-ATACTAGTGCTAGCTCTAGACTCGAGTATGGTGAG-3’ and reverse primer 5’-ATAGCGGCCGCTGATCAGCGGTTTAAACTTAAGC-3’. Next, the product and pDRF1-GW were digested using SpeI and NotI (New England Biolabs, Ipswich, Massachusetts, USA) and the PCR product was ligated into pDRF1-GW using T4 ligase (New England Biolabs). Next, a PCR was performed on tdTomato pDRF1-GW using FW primer 5’-ATGAATTCATGGTGAGCAAGGGC-3’ and RV primer 5’-ATGCGGCCGCTTACTTGTACAGCTCGTCCA-3’. The PCR product and the new AT1.03 and AT1.03^R122KR126K^ in pDRF1-GW were digested with EcoRI (New England Biolabs) and NotI. Afterwards, the PCR product was ligated into AT1.03 and AT1.03^R122KR126K^ in pDRF1-GW using T4 ligase, which replaced cp173-mVenus with tdTomato. Next, a PCR using KOD polymerase was performed on ymTq2 pDRF1-GW with FW primer 5’-CTGCTAGCACTAGTAAGCTTTTAA-3’ and RV primer 5’-ATATCGATAGCAGCAGTAACGAATTCC-3’ which produced ymTq2 with the last 11 amino acids removed (ymTq2Δ11). The PCR product and AT1.03 and AT1.03^R122KR126K^ in pDRF1-GW in pDRF1-GW were digested with NheI and ClaI (New England Biolabs). Next, the PCR product was ligated into AT1.03 and AT1.03^R122KR126K^ in pDRF1-GW using T4 ligase, which produced AT1.03^ymTq2Δ11-tdTomato^ and AT1.03^R122KR126K-ymTq2Δ11-tdTomato^, named yAT1.03 and yAT1.03^R122KR126K^.

### Yeast transformation

The yeast strain W303-1A (MATa, leu2-3/112, ura3-1, trp1-1, his3-11/15, ade2-1, can1-100) was transformed as described by Gietz and Schiestl.^31^

### ConA solution

Concanavalin A was prepared as described by Hansen et al., 2015.^32^ Briefly, 5 mg of Concanavalin A (Type IV, Sigma Aldrich) was dissolved in 5mL PBS at pH6.5, 40 mL H_2_O, 2.5mL of 1 M MnCl_2_ and 2.5 mL of 1 M CaCl_2_. This solution was aliquoted, snap-frozen and stored at -80°C.

### In vitro characterisation

W303-1A WT cells expressing yAT1.03 pDRF-GW and the empty pDRF1-GW vector were grown overnight at 200 rpm and 30 °C in 1x yeast nitrogen base without amino acids (YNB, Sigma Aldrich, Stl. Louis, MO, USA), containing 100 mM glucose (Boom BV, Meppel, Netherlands), 20 mg/L adenine hemisulfate (Sigma-Aldrich), 20 mg/L L-tryptophan (Sigma-Aldrich), 20 mg/L L-histidine (Sigma Aldrich) and 60 mg/L L-leucine (SERVA Electrophoresis GmbH, Heidelberg, Germany). Next, cells were diluted in 50 mL medium and grown to an OD_600_ of approximately 3. Cells were kept on ice and washed twice with 20 mL 0.01 M KH_2_PO_4_/K_2_HPO_4_ buffer at pH7 containing 0.75 g/L EDTA (AppliChem GmbH, Darmstadt, Germany). Next, cells were resuspended in 2 mL of 0.01 M KH_2_PO_4_/K_2_HPO_4_ buffer containing 0.75 g/L EDTA and washed twice in 1 mL of ice-cold 0.1M KH_2_PO_4_/K_2_HPO_4_ buffer at pH7.4 containing 0.4 g/L MgCl_2_ (Sigma-Aldrich). Cells were transferred to screw cap tubes containing 0.75 grams of glass beads (425-600 µm) and lysed using a FastPrep-24 5G (MP Biomedicals, Santa Ana, CA, USA) with 8 bursts of 6 m/s and 10 second. Lastly, the lysates were centrifuged for 15 minutes at 21000 g and the cell-free extracts were snap-frozen in liquid nitrogen.

Per sample, 5 wells of a black 96-well microtitre plate (Greiner Bio-One) were filled with 4 µL cell-free extract (10x diluted in distilled water) and 36 µL 10 mM HEPES-KOH buffer (Sigma-Aldrich) at various pH. Fluorescence spectra were recorded after subsequent additions of ATP (Sigma-Aldrich) using a CLARIOstar platereader (BMG labtech, Ortenberg, Germany). Spectra were obtained using 430/20 nm excitation and 460-660 nm emission (10 nm bandwidth). Fluorescence spectra were corrected for background fluorescence (by correcting for fluorescence of cell-free extract of cells expressing the empty pDRF1-GW plasmid) and FRET ratios were calculated by dividing acceptor over donor fluorescence. The dose-response curve was fitted to equation 1^23^, with FRET_max_ and FRET_min_ denoting the maximal and minimal FRET values obtained, ATP the ATP concentration (mM), k_d_ the dissociation constant (mM) and n the Hill coefficient (n).

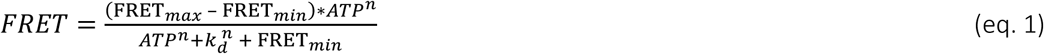

### Microscopy

Cells expressing yAT1.03 or yAT1.03^R122KR126K^ were grown overnight at 200 rpm and 30 °C in 1x YNB medium, containing 20 mg/L adenine hemisulfate, 20 mg/L L-tryptophan, 20 mg/L L-histidine, 60 mg/L L-leucine and either 1% ethanol (v/v, VWR International, Radnor, PA, United States of America), 100 mM fructose (Sigma Aldrich) or 111 mM galactose (Sigma Aldrich). Next, cells were diluted in the same medium and grown overnight to a maximum OD_600_ of 1.5 (midlog). Afterwards, the cells were transferred to a six-wells plate containing ConA coated coverslips. Coverslips with the attached cells were put in an Attofluor cell chamber (Thermofisher Scientific, Waltham, MA, USA) and 1 mL of fresh medium was added. Next, the coverslips were imaged using a Nikon Ti-eclipse widefield fluorescence microscope (Nikon, Minato, Tokio, Japan) at 30°C equipped with a TuCam system (Andor, Belfast, Northern Ireland) containing 2 Andor Zyla 5.5 sCMOS Cameras (Andor) and a SOLA 6-LCR-SB power source (Lumencor, Beaverton, OR, USA). FRET was recorded using a 438/24 nm excitation filter, a 483/32 nm donor emission filter and a 593/40 nm acceptor emission filter with a 552 nm long-pass (LP) dichroic filter (all filters from Semrock, Lake Forest, IL, USA). After recording the baseline, 111 µL of 1x YNB containing the necessary amino acids and a 10x amount of the desired substrate was added. For the multiple glucose pulses, subsequent additions of 20, 20, 40 and 80 µL of 50 mM glucose at 3, 10, 17 and 24 minutes were added to the cell chamber. At least 2 biological replicates were obtained for each experiment. Cells were segmented by an in-house macro using FiJi (NIH, Bethesda, MD, USA) and moving or dead cells were manually removed.

### Growth experiments

Cells expressing yAT1.03 and the empty pDRF1-GW vector were grown to midlog as described for microscopy with medium containing 1% ethanol. Cells were washed and resuspended to an OD_600_ of 1 with the same medium without any carbon source. Next, 20 µL of cells were transferred to a 48-wells plate with each well containing 480 µL of fresh medium with either 0.1% ethanol, 10 mM galactose or 10 mM glucose. Afterwards, cells were grown in a Clariostar plate reader at 30°C and 700 rpm orbital shaking. OD_600_ was measured every 5 minutes.

### Data analysis

R version 3.5.1 (R Foundation for Statistical Computing, Vienna, Austria) was used to analyse and visualize the obtained data. In brief, mTq2 bleedtrough was corrected and cells with a fluorescence below 500 counts (arbitrary units) were deleted. FRET ratio normalization was performed by dividing all FRET values by the mean FRET value of the baseline (before perturbations). Maximal FRET decrease and increase were determined with a sliding window of 3 frames. Cluster amounts were determined by using the fviz_nbclust function from the factoextra package.

## Results

### yAT1.03 has improved pH robustness

Experiments performed with AT1.03 showed severe drifts in FRET signal in unperturbed cells using our setup (Fig. S1). Furthermore, the original paper that reported AT1.03 already showed the pH sensitivity in the range of physiological pH of yeast^23^. We hypothesised that the baseline drift and the pH sensitivity are caused by the fluorescent proteins, since the donor mseCFP is poorly characterised and mVenus is known to be pH sensitive and not photostable^25^. Based on this, we undertook improvement of AT1.03 by changing the fluorescent proteins to ymTq2Δ11 (donor) and tdTomato (acceptor) as a more reliable and pH-insensitive FRET pair (Botman et al., in submission^33^). Replacements of mseCFP for ymTq2Δ11 and cp173-mVenus for tdTomato resulted in yAT1.03. *In vitro* characterisation showed a huge improvement in pH sensitivity compared to the original AT1.03 sensor^23^. With the new FPs, FRET ratios, at a fixed ATP concentration, remained stable across the normal physiological pH range (pH 6.25 – 7.25) of yeast (Fig. 2), indicating that the pH sensitivity of the original AT1.03 sensor indeed arose from the fluorescent proteins. The yAT1.03 sensor showed a K_d_ of 3.2 mM for ATP, which is almost identical to the original AT1.03. Lastly, expression of yAT1.03 in W303-1A had no effect on growth (Fig. S2), indicating that the sensor can be used without adverse effects on yeast physiology.

**Figure 2.**
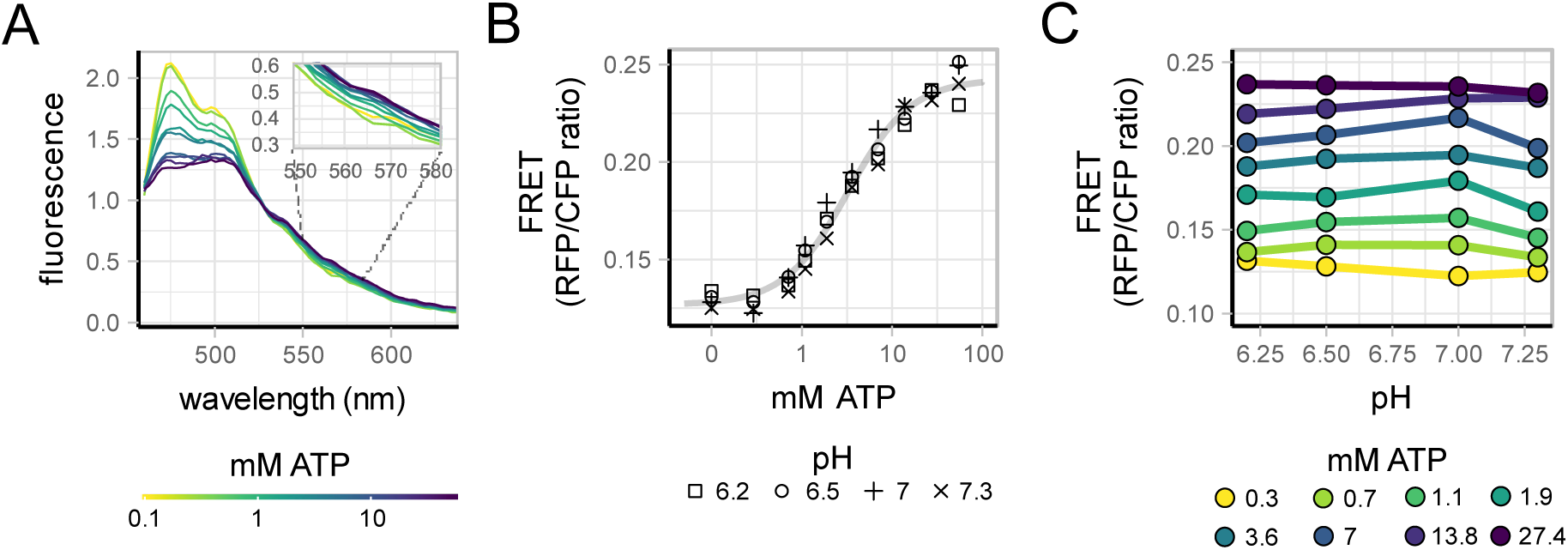
*In vitro* characterisation of yAT1.03. A) Fluorescence spectra of yAT1.03, obtained from cell-free extracts, gradient colour indicates ATP concentration. Inset shows fluorescence values in the acceptor range B) Dose-response curve of yAT1.03 obtained from the spectra, at various pH. Points indicate mean FRET ratio of 5 replicates, point shapes indicate pH C) pH stability of the sensor at various ATP concentrations, points indicate mean FRET ratio of 5 replicates, colours indicate the ATP concentration.

### yAT1.03 measures ATP reliably

*In vitro* characterisation of the sensor showed robust ATP responses of the sensor. To verify that yAT1.03 visualizes ATP changes reliably *in vivo* as well, we performed several control experiments (Fig. 3). First, we tested whether the sensor showed a typical transient change in ATP when cells experience a sudden glucose perturbation (Fig. 3A). As expected, yAT1.03 FRET ratios transiently decreased, followed by a recovery, while no response was observed when medium without glucose was pulsed. -The same glucose perturbation did not elicit a response in the non-responsive sensor yAT1.03^R122KR126K^. These results imply that the sensor indeed measures ATP and no other effects.

**Figure 3.**
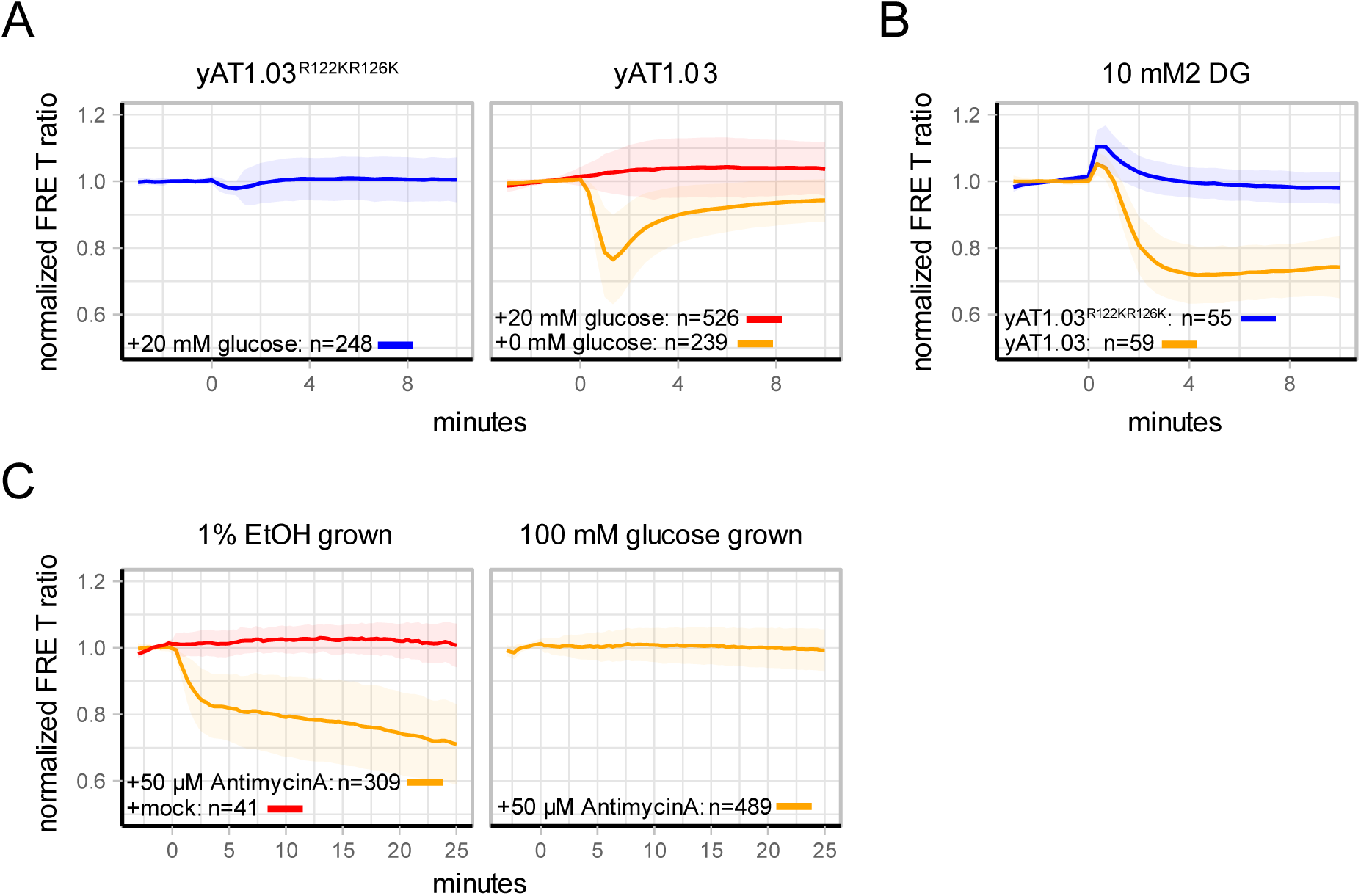
*In vivo* experiments show reliable yAT1.03 output. A) W303-1A WT cells expressing yAT1.03 or yAT1.03^R122KR126K^ (depicted above each graph) were grown on 1% EtOH. At t=0 minutes, glucose or the same medium without glucose was added and the FRET responses were measured. B) W303-1A WT cells expressing yAT1.03 or yAT1.03^R122KR126K^ were grown with 1% EtOH and incubated in 10 mM glucose for at least 1 hour. Afterwards, cells were visualized and 10 mM 2-DG was added at t=0 minutes. C) W303-1A WT cells expressing yAT1.0 were grown with 1% EtOH or 100 mM glucose as substrate (depicted above each graph). Antimycin A or only the solvent (mock) was added to the cells at t=0 minutes. Lines show mean responses, normalized to the baseline, shaded areas indicate SD, color indicates either the sensor expressed or the added solution. Percentages are v/v, abbreviations: EtOH, ethanol

Next, we tested if an extreme perturbation of metabolism affects yAT1.03 readouts (Fig. 3B). Cells were incubated for 60-90 minutes in 10 mM glucose after which the glycolytic inhibitor 2-deoxy-D-glucose (2-DG) was added to the cells. 2-DG is transported and phosphorylated by ATP, but not further metabolised, and thus acts as an ATP drain^34,35^. FRET responses showed a rapid decrease of FRET, which confirms that the sensor faithfully reports depletion of the ATP pool caused by 2-DG. In contrast, the non-responsive sensor yAT1.03^R122KR126K^ showed only a minor response, indicating that the yAT1.03 sensor performs robustly in response to extreme metabolic perturbations. Lastly, we tested whether we could distinguish whether ATP is generated by respirative or fermentative metabolism (Fig. 3C). Cells were grown with either 1% ethanol as substrate (ATP generation entirely dependent on respiration) or 100 mM glucose as substrate (ATP generation largely through fermentation). Subsequently, 50 µM of antimycin A was added to block respiration through inhibition of the mitochondrial electron transport chain complex III. As anticipated, addition of antimycin A only depleted ATP levels when cells were growing on ethanol as a substrate. In conclusion, these results demonstrate that yAT1.03 can be used for robust measurements of ATP dynamics *in vivo*.

### Pre-growth conditions largely determine ATP responses during transitions

After establishing yAT1.03 as a robust ATP sensor, we used it to characterise ATP dynamics in response to different carbon source transitions. Cells were grown on various carbon sources and transitioned to glucose (the preferred carbon source) or galactose (Fig. 4A). Cells grown on fructose showed only a small transient ATP response when challenged 100 mM glucose, and no response with 20 mM glucose. In contrast, glucose addition to galactose grown cells induced the biggest transient decrease of ATP with a mean decrease of 37% in FRET. Lastly, glucose and galactose addition to ethanol-grown cells both show a response, but the responses are qualitatively very different. Glucose addition results in a transient FRET decrease of 24%. In contrast, galactose addition lacked a transient ATP response, but showed a steady decrease to the same level compared to glucose, 10 minutes after addition. Interestingly, both the decrease and recovery of ATP show a high degree of variability between cells, indicating heterogeneity in the responses of individual cells (Fig. 4A,B).

**Figure 4.**
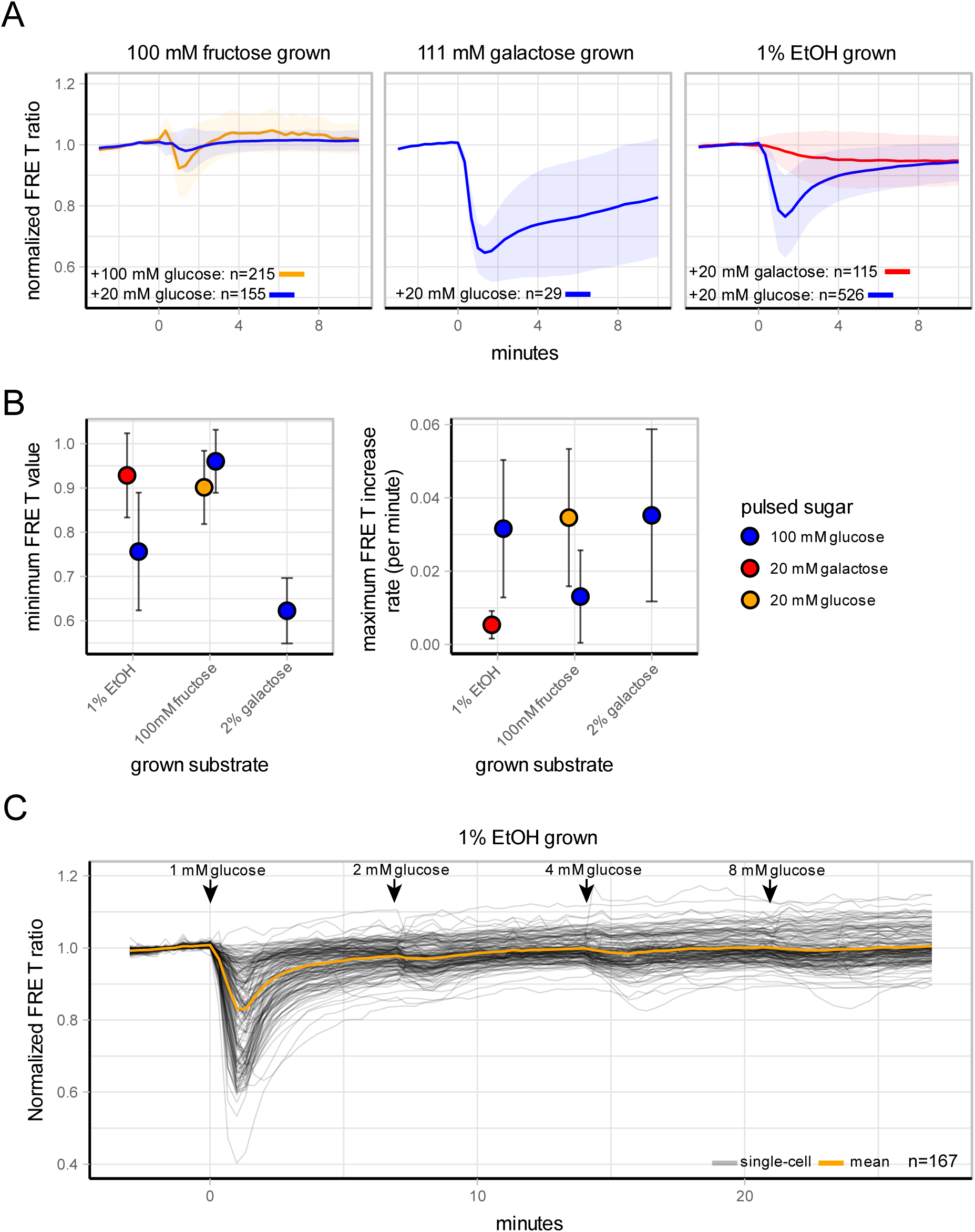
ATP dynamics during various carbon transitions. A) ATP dynamics of W303-1A cells expressing yAT1.03 grown on either 1% EtOH, 100 mM fructose or 111 mM galactose and pulsed with either 20 mM glucose, 100 mM glucose or 100 mM galactose. Lines show mean FRET ratios, normalized to the baseline, shade areas indicate SD. B) Minimum FRET value (i.e. maximum decrease of ATP levels) and ATP recovery speed (depicted by maximum FRET increase per minute) of each transition. Points indicate mean value, error bars indicate SD. C) ATP dynamics of W303-1A cells expressing yAT1.03 grown on 1% EtOH and pulsed successively with increasing amounts of glucose. Arrows indicate time points of glucose addition. Orange line show mean FRET ratio (baseline normalized, grey lines show single-cell traces.

Finally, since we were able to conveniently measure ATP in time in living cells, we looked at how ATP levels change in response to multiple successive glucose additions (Fig. 4D). The first addition of only 1 mM of glucose to ethanol-growing cells induced a clear transient ATP decrease but subsequent additions showed only slight responses for most cells. Still, we found again heterogeneity in the responses of individual cells as can be seen from single-cell traces in figure 4 and S4.

In summary, we show that ATP dynamics are dependent on the pre-growth condition and show large variations among individual cells. Moreover, for most cells, 1 mM of glucose is sufficient to diminish subsequent ATP responses to further sudden increases in glucose.

### ATP dynamics during glycolysis start-up are heterogeneous between cells

Since the ATP responses showed high variability, we looked in more detail at single-cell responses during the ethanol to 20 mM glucose transition (Fig. 5). Visualization of the single-cell trajectories and their distributions clearly showed large heterogeneity during the response (Fig. 5A,B and movie S1). We used a hierarchical clustering-approach (using Euclidean clustering) to see if dynamic traces can be grouped into distinct response-classes (Fig. 5C). Next, we determined the optimal cluster amounts (3) using the silhouette method (Fig. S3). We identified three types of responses, and determined various characteristics for each, such as the absolute baseline FRET ratio (the not-normalized FRET ratio before sugar addition), change in the normalized FRET values at the end of the timecourse compared to the baseline (ΔFRET), the maximal FRET decrease- (ATP depletion rate) and increase rate (ATP recovery rate), the minimum FRET value (lowest amount of ATP obtained after the transition) and the time to reach this minimum FRET value (Fig. 5D). No difference in the non-normalized baseline FRET values could be found between the clusters, indicating that the starting ATP alone does not determine the ATP dynamics after a transition to glucose. The first cluster contained approximately 61% of all responses and showed a small transient decrease in ATP followed by rapid recovery to pre-perturbation levels after approximately 8 min. The second cluster contained 36% of the cells and showed a large transient decrease in ATP, followed by fast, but not yet full, recovery after 12 min. A third, smaller cluster, contained 3% of all cells and was characterized by rapid ATP depletion, similar to perturbations with either 2-DG or antimycin A (Fig. 2), with very slow recovery (or even absent for some cells in this cluster). The absence of ATP recovery in some cells is indicative of an imbalanced metabolic state that cells can get trapped in during these transitions^17^. Such heterogeneous ATP responses were also found when cells were pulsed successively with increasing amounts of glucose (Fig. S4). Here, some cells only showed an ATP response after the first glucose pulse, some cells show a response after each addition, while yet others show no responses to any of the perturbations. In contrast to dynamic conditions, steady-state ATP levels of glucose-grown cells showed no clear subpopulations (Fig. S6). In summary, we show that glycolytic start-up is highly heterogeneous, ranging in the one extreme from cells that show small or no transient changes in ATP, to cells that end up in an imbalanced state, with no or little recovery of ATP after a glucose pulse (on short time-scale).

**Figure 5.**
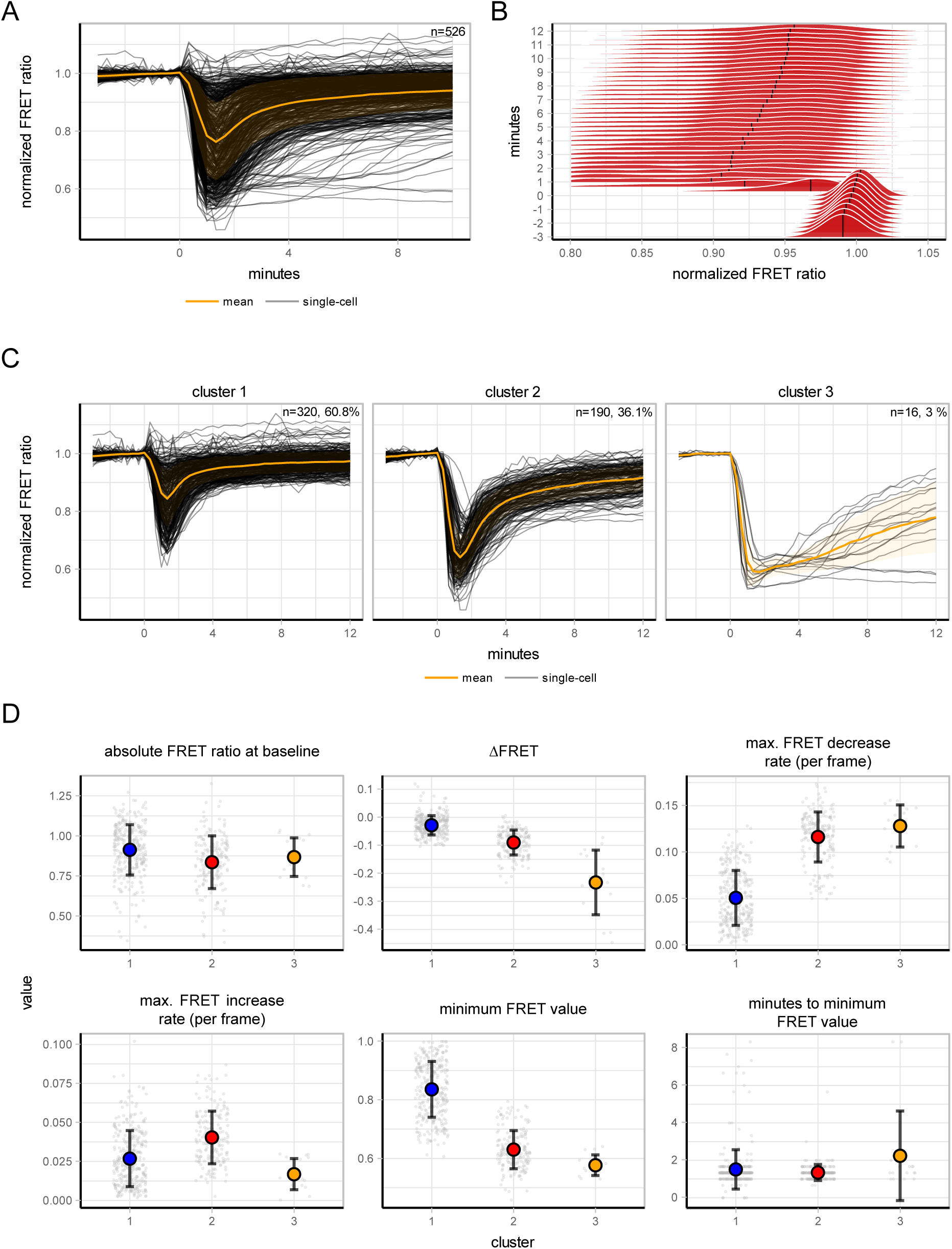
Heterogeneity of cells transitioned from 1% EtOH to 20 mM glucose. A) ATP dynamics of W303-1A cells expressing yA1.03 grown on either 1% EtOH and pulsed with 20 mM glucose. Orange line shows mean normalized FRET ratios, shades indicate SD, grey lines show single-cell traces. B) Normalized frequency distributions of the same transition depicted by graph A. C) Hierarchical clustering of the single-cell trajectories obtained from graph A showed 3 distinct subpopulations. Orange lines show normalized mean FRET ratios, shades indicate SD, grey lines show single-cell traces D) Various ATP dynamic parameters per cluster. Absolute FRET ratio at baseline depict the mean FRET value (not normalized) of the first 10 frames (before glucose addition), ΔFRET is calculated as the difference of the mean FRET value of the last 5 frames compared to the baseline. Dots indicate single-cell values, points indicate mean values for each cluster, errorbars indicate SD.

## Discussion

Previous work showed that nutrient transitions can result in phenotypic subpopulations, and suggested that these subpopulations are a consequence of non-genetic metabolic difference between individual cells^17^. We therefore require tools to study metabolic dynamics at the single cell level. To gain insights into the dynamics of ATP in single yeast cells, we adapted the AT1.03 sensor to make it more suitable for readouts under dynamic conditions. Specifically, metabolic dynamics in yeast are often characterized by pH changes^17,27^, and the AT1.03 sensor is sensitive in the physiological pH-range of *S. cerevisiae*^23^. Also, ATP and pH homeostasis are intimately coupled and this presents an obvious challenge: simultaneous changes in ATP and pH will significantly confound sensor-signal and its interpretation. Our modified sensor shows improved pH stability in the required physiological range (pH 6.25 – 7.25) and has a K_d_ of 3.2 mM, which is optimal given intracellular ATP concentrations in yeast between 1-4 mM^14– 17,36–39^. Our control experiments show that the sensor can be used to reliably measure ATP levels, even for rather extreme metabolic perturbations. First, treatment of fermenting cells with the glycolytic inhibitor 2-DG showed clear ATP drainage. Second, inhibition of the electron transport chain only affects ATP levels in cells growing on ethanol (respiring), and not on glucose (fermenting). In addition, behaviour of the non-responsive yAT1.03^R122KR126K^ demonstrated specificity towards ATP.

We characterized ATP dynamics in response to various sudden carbon source transitions as a readout for glycolytic activation. We found that ATP-response depended significantly on the combination of pre-growth conditions and pulsed carbon source. For example, galactose-grown cells have a bigger ATP depletion compared to ethanol grown cells, indicating a large imbalance between ATP consumption and production during start-up of glycolysis (Fig. 3). This suggests that cells grown on galactose may have a higher glucose-phosphorylation capacity in the upper-part of glycolysis compared to ethanol, which is in line with the fact that galactose is a glycolytic substrate, and ethanol a gluconeogenetic one. Indeed, previous reports state that galactose-grown cells have a higher expression of the hexokinases and glucose transport capacity^15,40–42^. In addition, we show that galactose addition to ethanol grown cells does not cause a rapid transient decrease in ATP, but rather a gradual reduction in ATP levels towards a new steady-steady state. These differences indicate that galactose does not cause the same transient imbalance in ATP consumption and production that glucose does when cells are pre-cultured on ethanol. Although the first step in galactose metabolism also involves substrate phosphorylation, and therefore ATP consumption, this activity will initially be low as the galactose metabolising Leloir-pathway is only fully induced in the presence of galactose. We also measured ATP dynamics in fructose grown cells subjected to sudden glucose additions. The small or absent response to glucose indicates that glycolysis is hardly perturbed. This makes sense, as based on growth rates, the glycolytic flux is similar for these sugars. These results combined indicate that the transient ATP decrease is caused by an imbalance between upper (ATP consuming) and lower (ATP producing) parts during the start-up of glycolysis. The timing at which cells experience their minimal ATP levels (minutes of maximal dip) is independent of the extent of ATP decrease (Figs. 4A and S5). This suggests that a larger initial ATP depletion is not caused by a longer duration of the upper and lower-glycolytic imbalance, but by the magnitude of the imbalance.

Lastly, we pulsed cells sequentially with increasing amounts of glucose (Fig. 4C). Most cells display a transient decrease of ATP only in response to the first addition of 1 mM of glucose. After that, the majority of the cells display no (or diminished) ATP-response to successive glucose additions. Ethanol-grown cells express HXT6 and HXT7 with a K_m_ around 1 mM^41^. Apparently, the addition of extra glucose to these cells does not cause a new imbalance between the upper and lower glycolysis.

Our clustering analysis of the ethanol to 20 mM glucose transition (Fig. 5) showed that a small fraction of approximately 3% of cells showed this phenotype. An earlier study showed that sudden transitions to glucose cause small populations of cells to end up in a low pH state, a state inferred to indicate an upper- and lower glycolytic imbalance^17^. Our ATP measurements support this inference, showing that indeed a small subpopulation of cells have trouble balancing ATP consumption and production during glycolytic start-up. Previously, it was shown that small variations in metabolic parameters, including enzyme expression levels and metabolite concentrations determine how a cell will respond to sudden glucose addition. Our data suggests that differences in glycolytic start-up dynamics cannot be explained solely by initial ATP levels (Fig 5D).

In conclusion, we provide the yAT1.03 sensor, which shows more robust ATP measurements in yeast cells compared to the original AT1.03 sensor, and specifically under dynamic conditions. With this sensor we could show that ATP dynamics during glycolytic start-up depends on pre-growth conditions and that isogenic cells show highly heterogeneous responses during transitions to glucose. We believe that the yAT1.03 sensor will be a useful addition to the current arsenal of tools to investigate ATP physiology.

## Acknowledgements

We greatly thank Joachim Goedhart (Molecular Cytology, University of Amsterdam) and Daan de Groot for fruitful discussions. We also thank Joachim Goedhart for sharing the clustering script on GitHub (https://github.com/JoachimGoedhart).

## Material requests

Plasmids and sequences are available at Addgene (https://www.addgene.org/Bas_Teusink/).

## Supplements

Movie S1. Ratiometric movie of W303-1A WT cells expressing yAT1.03. Cells were grown on 100 mM pyruvate and pulsed with 100 mM glucose. Colour indicates the FRET ratio, shown by the calibration bar in the upper left.

**Figure S1.**
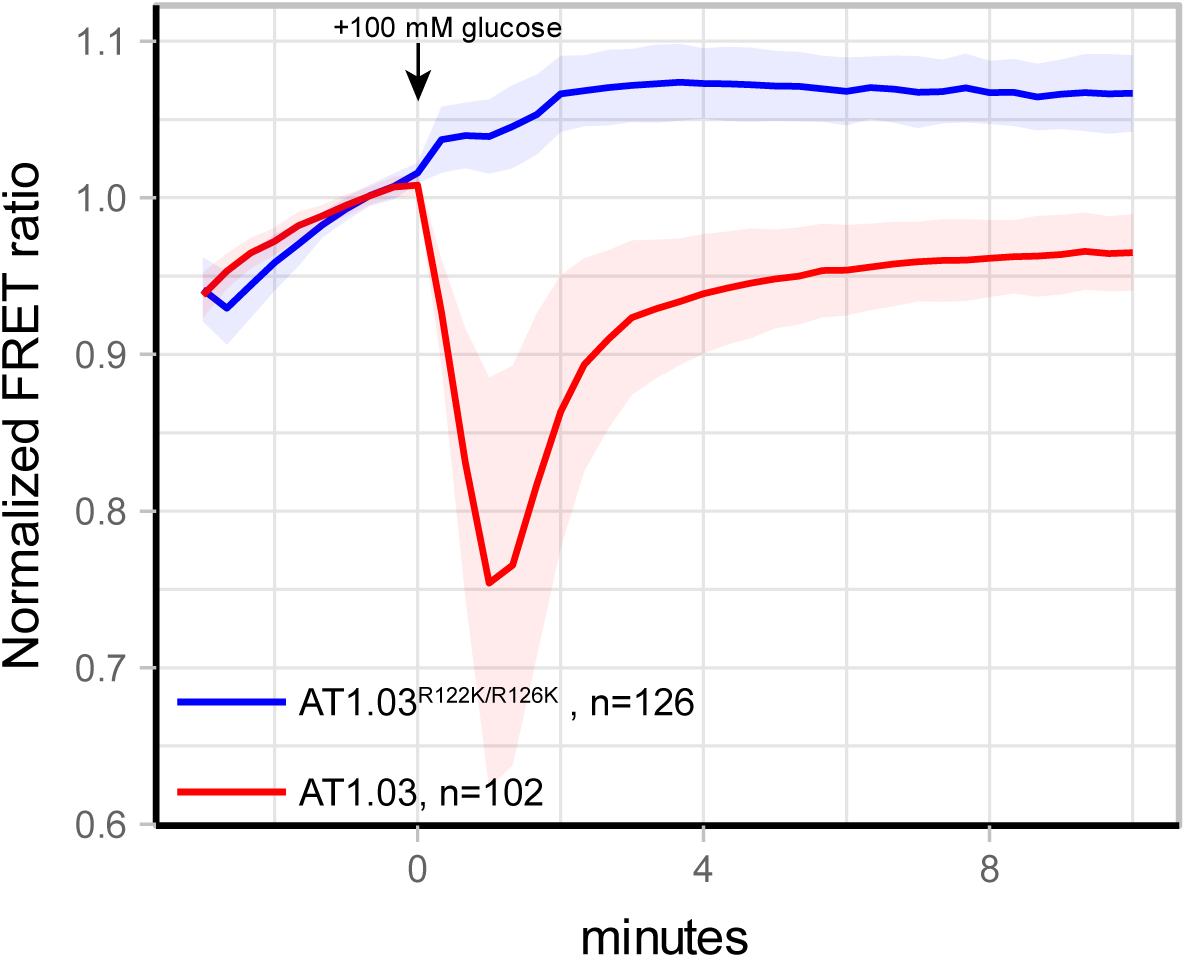
Baseline drift of the original AT1.03 sensor. W303-1A WT cells expressing yAT1.03 or yAT1.03^R122KR126K^ were grown with 1% EtOH as carbon source. At t=0 minutes, glucose was added and the FRET responses were measured. Lines show mean responses, normalized to the 5 last frames before glucose addition, shades indicate SD.

**Figure S2.**
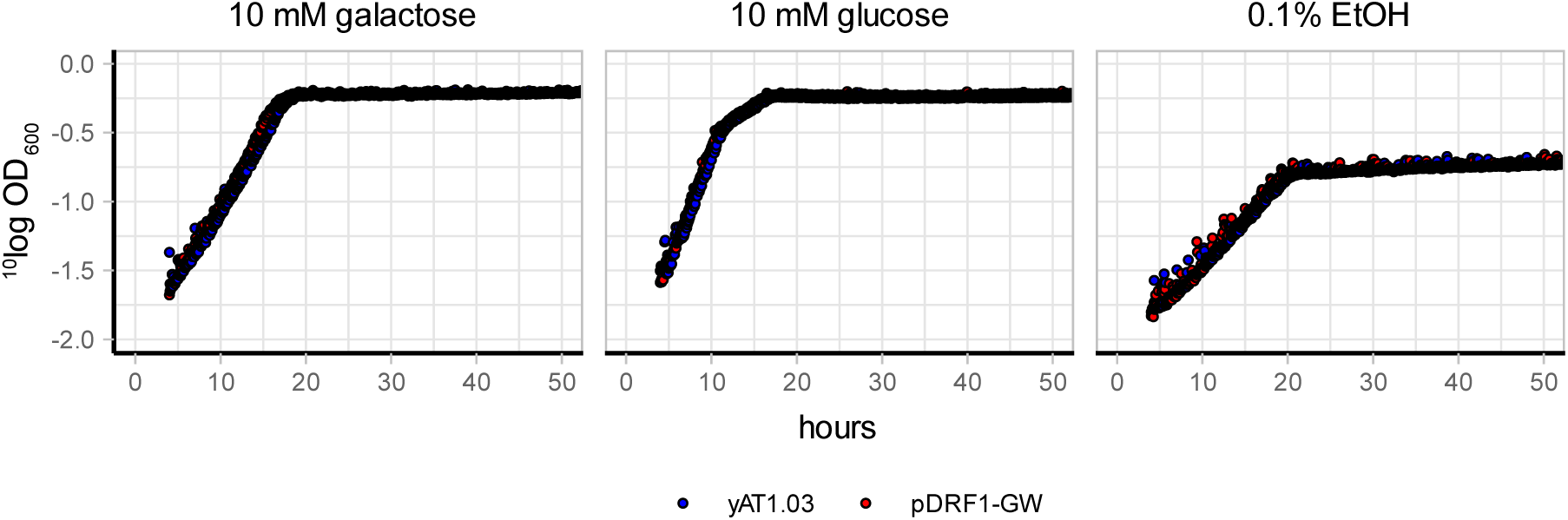
Growth of W303-1A cells expressing either yAT1.03 or the empty vector pDRF1-GW. Cells were grown to midlog with 0.1% EtOH as substrate. Next, cells were washed and incubated for 10 minutes in 1x YNB containing no carbon source. Next, cells were transferred to 1x YNB containing either 10 mM galactose, 10 mM glucose or 0.1% EtOH and OD_600_ was measured. Dots depict the ^10^log value of the OD, colours indicate the strain.

**Figure S3.**
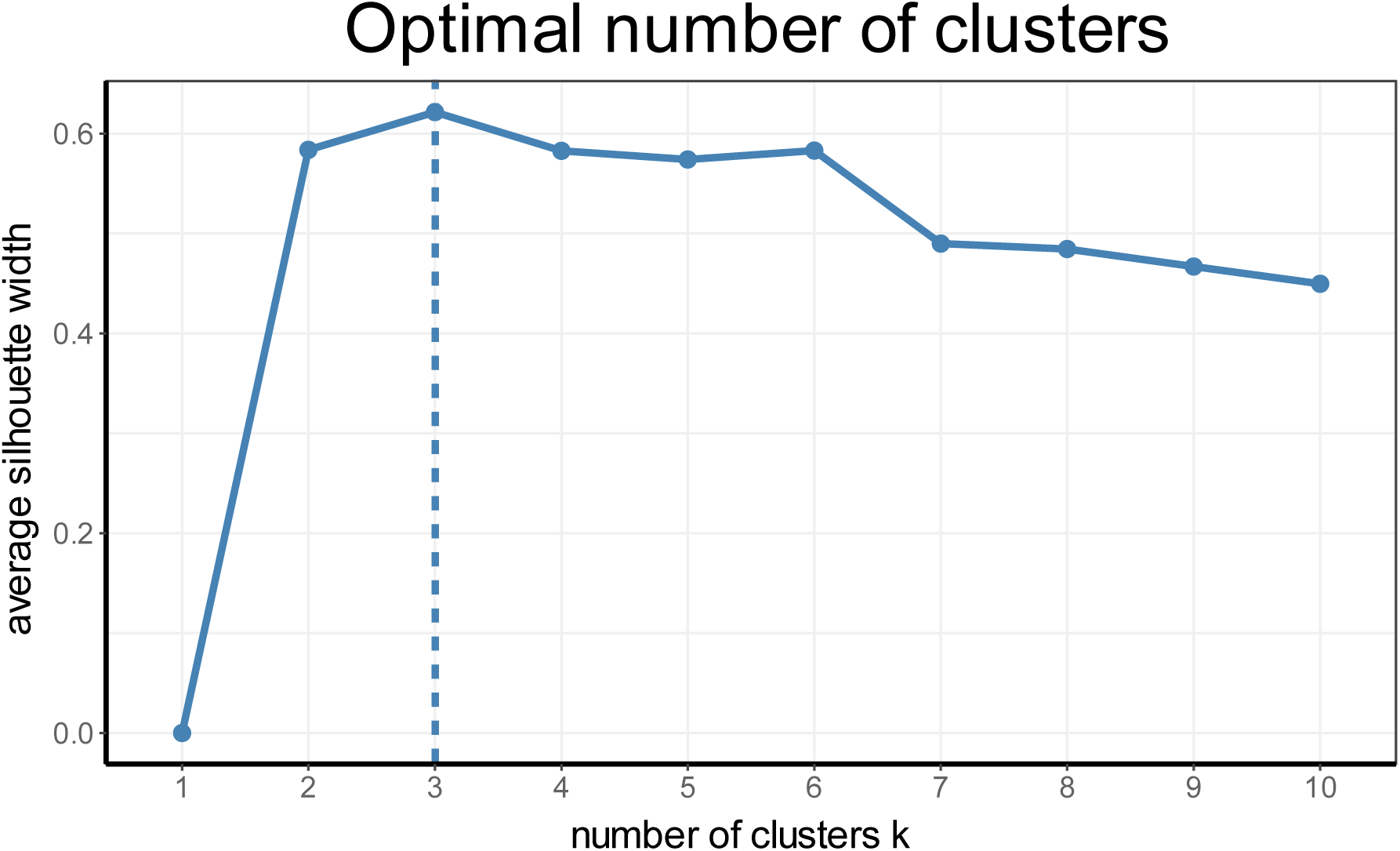
Determination of the optimal number of clusters. Three clusters were chosen based on the silhouette method, using the fviz_nbclust function in R.

**Figure S4.**
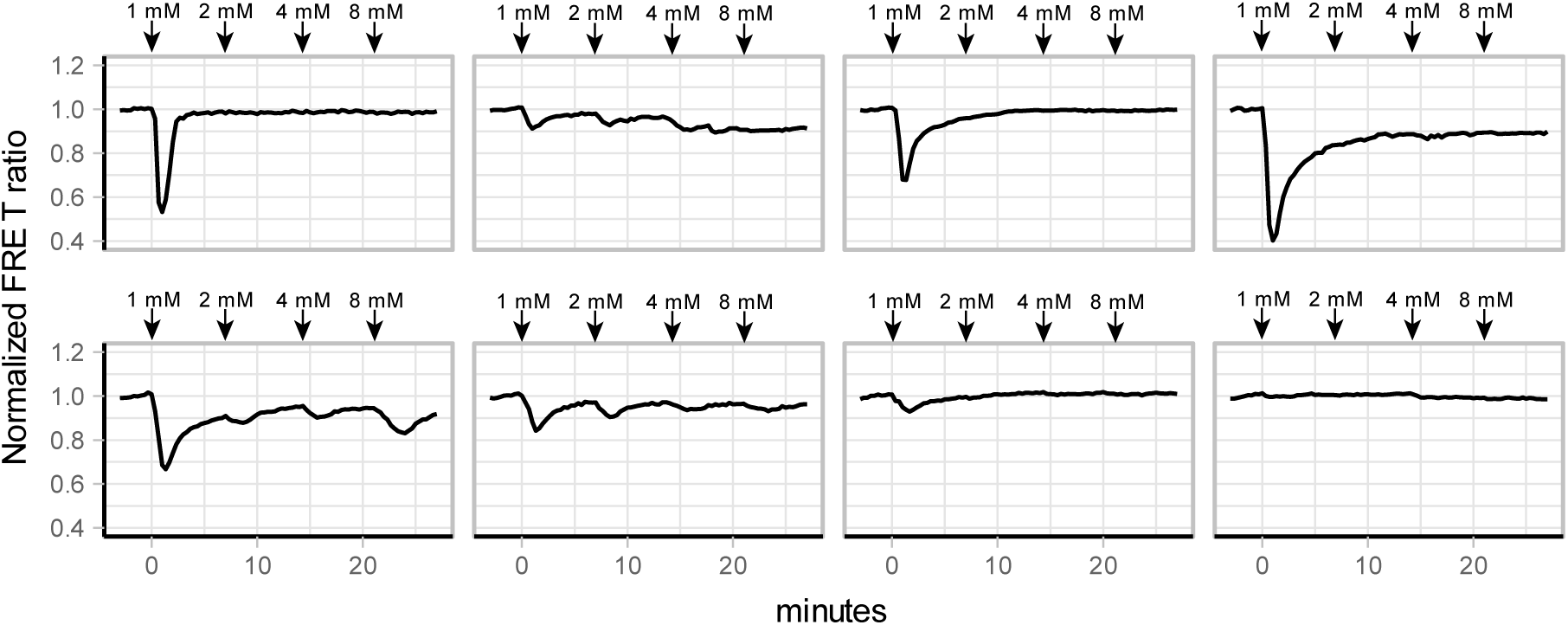
Single-cell FRET trajectories of cells exposed to multiple glucose pulses. W303-1A cells expressing yA1.03 were grown on 1% EtOH and pulsed multiple times with increasing amounts of glucose. Every graph depicts the ATP response of a single-cell. Arrows indicate point of the new glucose concentrations reached. Lines show single-cell traces, normalized to the baseline.

**Figure S5.**
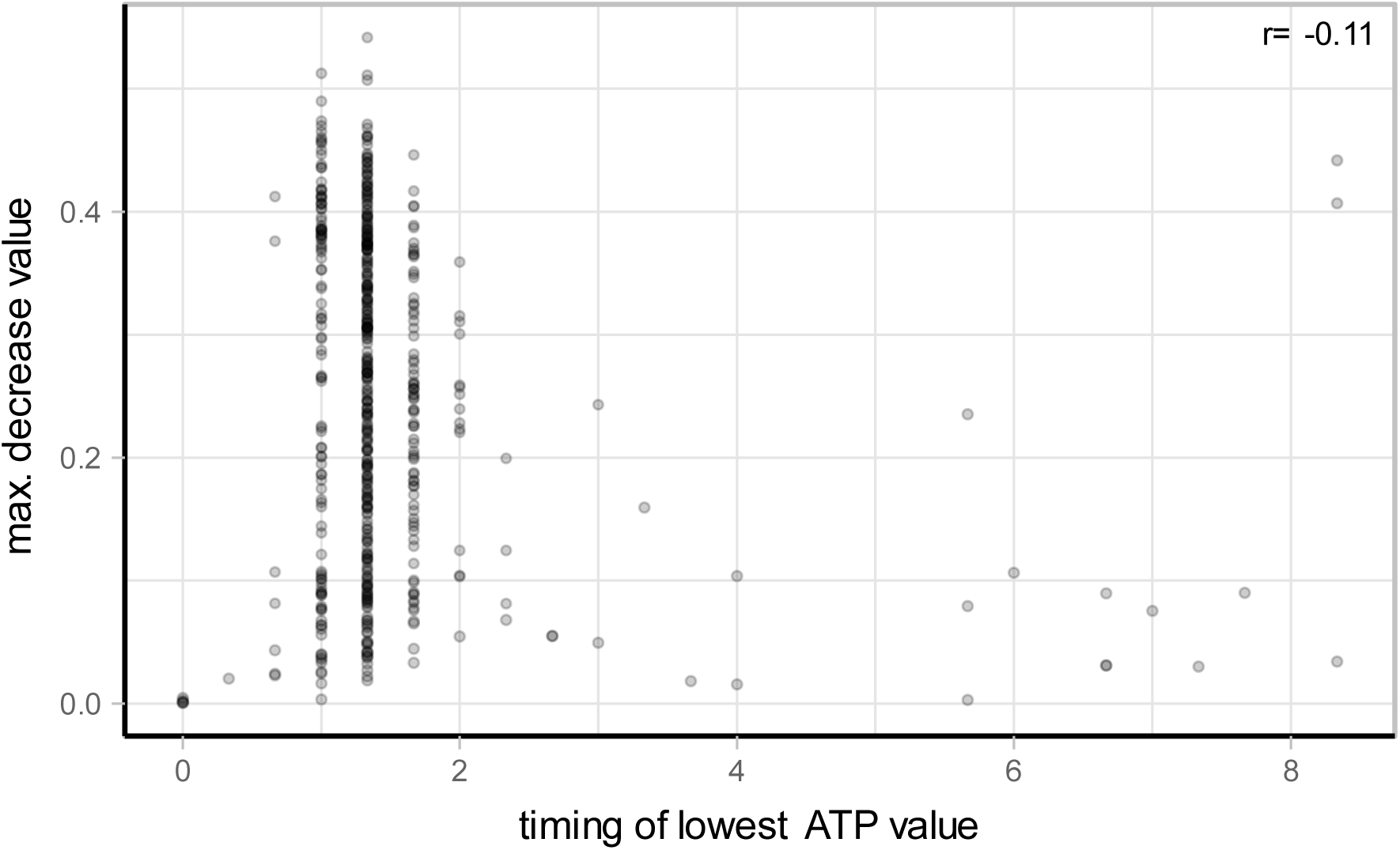
Correlation between the maximum decrease of ATP in a cell and the timing of this observed dip. Points show single-cell values. The spearman correlation coefficient (r).

**Figure S6.**
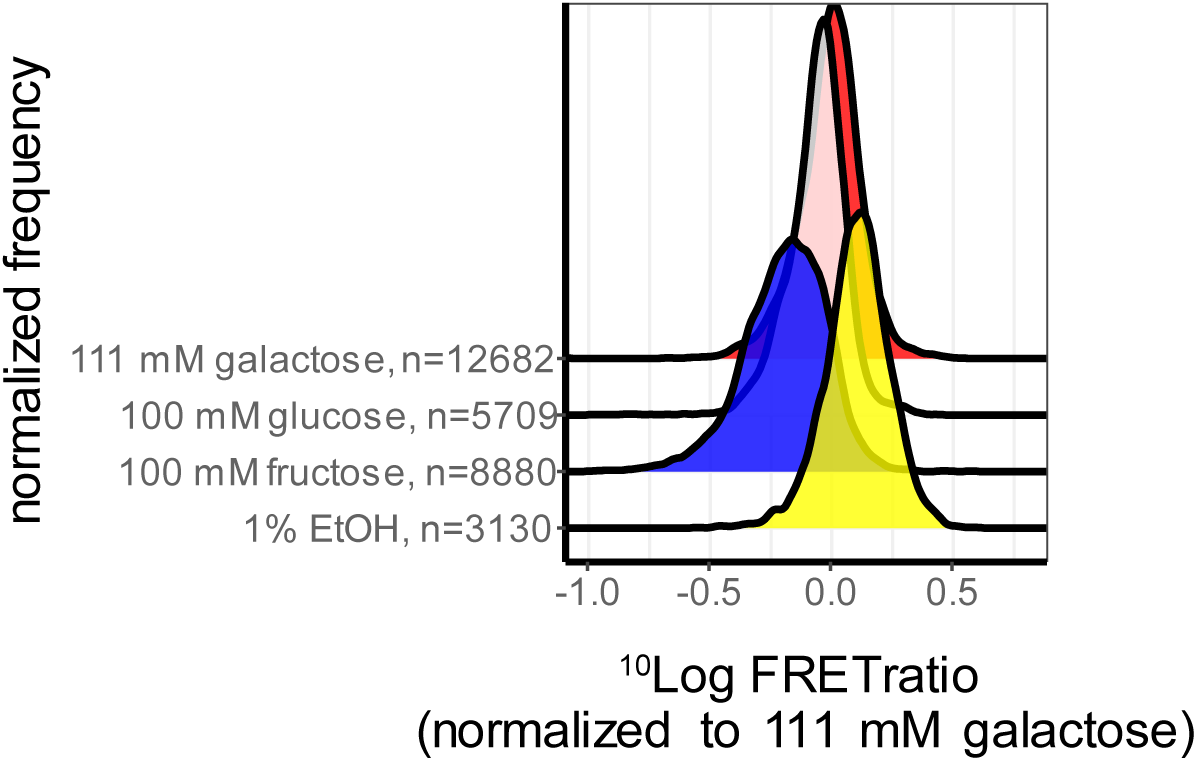
Steady-state ATP levels of W303-1A cells grown on various substrates. A) Normalized frequency distributions of W303-1A cells grown on 1% EtOH, 100 mM fructose, 100 mM glucose or 111 mM galactose. FRET values were normalized to cells grown on 111 mM galactose. B) Mean FRET ratios of W303-1A cells grown on the various substrate, normalized to 111 mM galactose. For every replicate, the median FRET ratio was determined and the mean of these values are indicated by the points. Point colour indicates the substrate grown on, errorbars indicate SD.

